# Modeling aroma release during *in vivo* food oral processing using digimouth

**DOI:** 10.64898/2026.10.01.755679

**Authors:** C. Peltier, O. Mourabit, K. Coulibaly, I. Andriot, I.C. Trelea

## Abstract

The perception of flavor during food consumption is a complex process governed by the dynamic release of aroma compounds, which is influenced by the interaction between the food matrix and human physiology. While mechanistic models have been developed to simulate these phenomena, their practical application has often been limited by the lack of flexible software tools capable of integrating diverse food structures and real-time physiological data. This study addresses this gap by incorporating *in vivo* mechanistic models for both liquid foods and melting solid matrices in a modular Python-based framework: digimouth.

The modeling approach treats the oral, pharyngeal, and nasal cavities as a series of interconnected reactors with mass transfer governed by ordinary differential equations (ODEs). These models explicitly integrate key physiological parameters, including salivary flow, bidirectional respiratory airflow, and discrete events such as mastication and swallowing. For solid foods, the framework incorporates additional mechanisms for matrix dissolution, temperature-dependent melting, and dynamic surface area change.

The models were validated using experimental data from 30 subjects who consumed aroma solutions and flavored gelatin gummy discs while monitored by PTR-MS (Proton Transfer Reaction – Mass Spectrometry). The simulations successfully reproduced the main kinetic features and release profiles observed experimentally for both types of matrices. Despite some limitations related to the self-reporting of physiological events and the non-inclusion of initial orthonasal mass transfer in the models, the developed software package proved to be a robust and evolutive tool. It offers a foundation for exploring inter-individual variability and the impact of complex food structures on sensory perception.

## 1. Introduction and context

The study of aroma release in food systems is a highly interdisciplinary field that combines principles of physical chemistry, mass transport phenomena, and dynamic process conditions. Initial modeling approaches were largely based on *in vitro* experiments designed to better understand the fundamental mechanisms controlling volatile compound release after a disruption of equilibrium, such as when a food product is exposed to air. These early investigations highlighted several key governing parameters, including air–product partitioning behavior and mass transfer coefficients in both the liquid and gas phases (Marin et al., 1999). Later, more advanced models have expanded classical mass transfer descriptions to encompass more structurally complex food systems, including emulsions and semi-solid matrices ( Harrison and Hills, 1997, Andriot et al., 2000). In such systems, interactions between aroma molecules and food matrix components can play a significant role in shaping release kinetics.

This *in vitro* foundational understanding paved the way for *in vivo* mechanistic compartmental models that simulate the real-time flow and concentration of aroma compounds within the oral, pharyngeal and nasal cavities, treating these anatomical spaces as interconnected reactors (Trelea et al., 2007; Doyennette et al., 2011). These ODE (Ordinary Differential Equation) models, pioneered by Trelea et al. (2007), and further developed by Doyennette et al., (2011, 2014), integrate key physiological factors—such as salivary flow rates, breathing patterns, and the volumes of anatomical compartments—to accurately predict aroma concentration over time. They highlighted the role of post-deglutitive pharyngeal residue, a thin liquid film left on the pharynx walls that continues to release aroma into the breath stream and which is subject to significant dilution by saliva. To address solid foods requiring mastication, these models evolved to incorporate the physical breakdown process (Doyennette et al., 2014) which mathematically parameterized fragmentation by linking chewing activity to a continuous dynamic increase in the solid/liquid contact surface area within the forming bolus. This approach also emphasized the continuous exchange of aroma-rich air between the oral cavity and the pharynx during chewing, allowed by the velopharynx opening. For specific food structures, such as cooked rice, a detailed conceptual model was proposed and validated *in vivo* by How et al. (2021), linking aroma release intensity directly to three main mechanistic pathways: the increase in particle surface area via a specific “cleave and paste” fragmentation process, the diluting effect of saliva, and the continuous diffusion of volatiles from residues trapped in the buccal pouches.

Although mechanistic *in vivo* models of aroma release provide strong conceptual and predictive capabilities, their practical use remains relatively limited. Recently, the authors developed a flexible Python package enabling the simulation of *in vitro* aroma release phenomena and facilitating the implementation of various modeling approaches (Peltier et al., 2027). Building upon this framework, the present study extends the package to incorporate *in vivo* aroma release models applicable to both liquid systems and melting solid matrices.

To evaluate the implementation, model predictions were generated and compared to experimental datasets obtained from mono-aroma model products, including aroma solutions and gelatin gummy discs (Andriot et al., 2026b). Simulated release profiles were subsequently confronted with experimental measurements in order to assess the ability of the models to reproduce real aroma release behavior.

## 2. Material and methods

### 2.1. Modeling *in vivo* aroma release in solutions

The *in vivo* model for solutions describes the retronasal transfer of volatile compounds during oral food consumption, from the oral cavity to the nasal cavity, in order to predict the concentration reaching olfactory receptors and measured by an analytic instrument (PTR-MS) connected to a device positioned at the nostrils of a subject. The model comprises six well-mixed compartments: the solution, the oral cavity air, the liquid film in the pharynx, the pharyngeal air, the nasal cavity, and the PTR-MS measurement compartment.

Compared to the *in vitro* setup (Peltier et al., 2027), two additional phenomena are explicitly considered: saliva flow, which affects dilution and transport, and breathing, which induces time-variant airflow and strongly influences aroma transport. Transport between compartments is governed by interfacial mass transfer, airflow induced by breathing and swallowing, and deposition of a liquid film after swallowing.

Additionally, two physiological events are incorporated: swallowing and jaw movements. Swallowing reduces the product volume in the oral cavity to a minimal value, deposits a thin film of product in the pharynx, and leads to complete mixing of all air compartments. Simulated chewing does not reduce the oral volume but opens the velopharynx, thereby mixing all air compartments and inducing aroma pulses in the nasal cavity.

#### 2.1.1. Model parameters

Figure 1 shows the different anatomic compartments, the state variables and the parameters included in the model to describe volatile mass transport when consuming a flavored solution.

**Figure 1:**
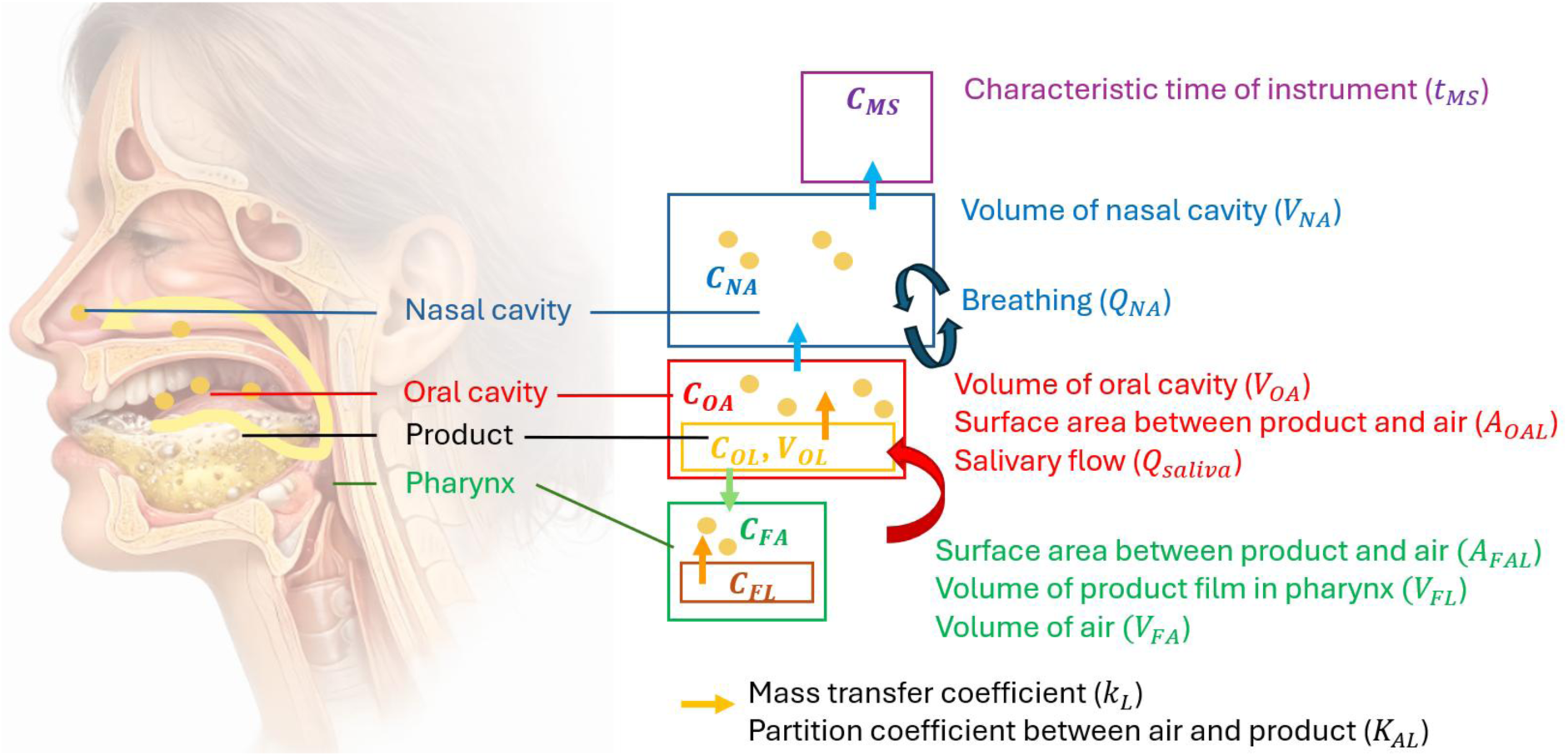
Illustration of the anatomic compartments, parameters and state variables of the model of solution tasting.

The state variables are in bold italic in the figure 1:

- *V_OL_*, the volume of liquid product (here, the aromatized solution) in the oral cavity,
- *C_OL_*, concentration of aroma in the liquid,
- *C_OA_*, concentration of aroma in the air contained in the oral cavity,
- *C_FA_*, concentration of aroma in the air contained in the pharynx,
- *C_FL_*, concentration of aroma in the product layer deposited in the pharynx,
- *C_NA_*, concentration in the nasal cavity,
- *C_MS_*, concentration measured by an instrumental device (here, PTR-MS, Le Quéré and Schoumacker, 2023).

The parameters are related to each compartment and initial values of each state variable. They are summarized in Table 1.

**Table 1:**
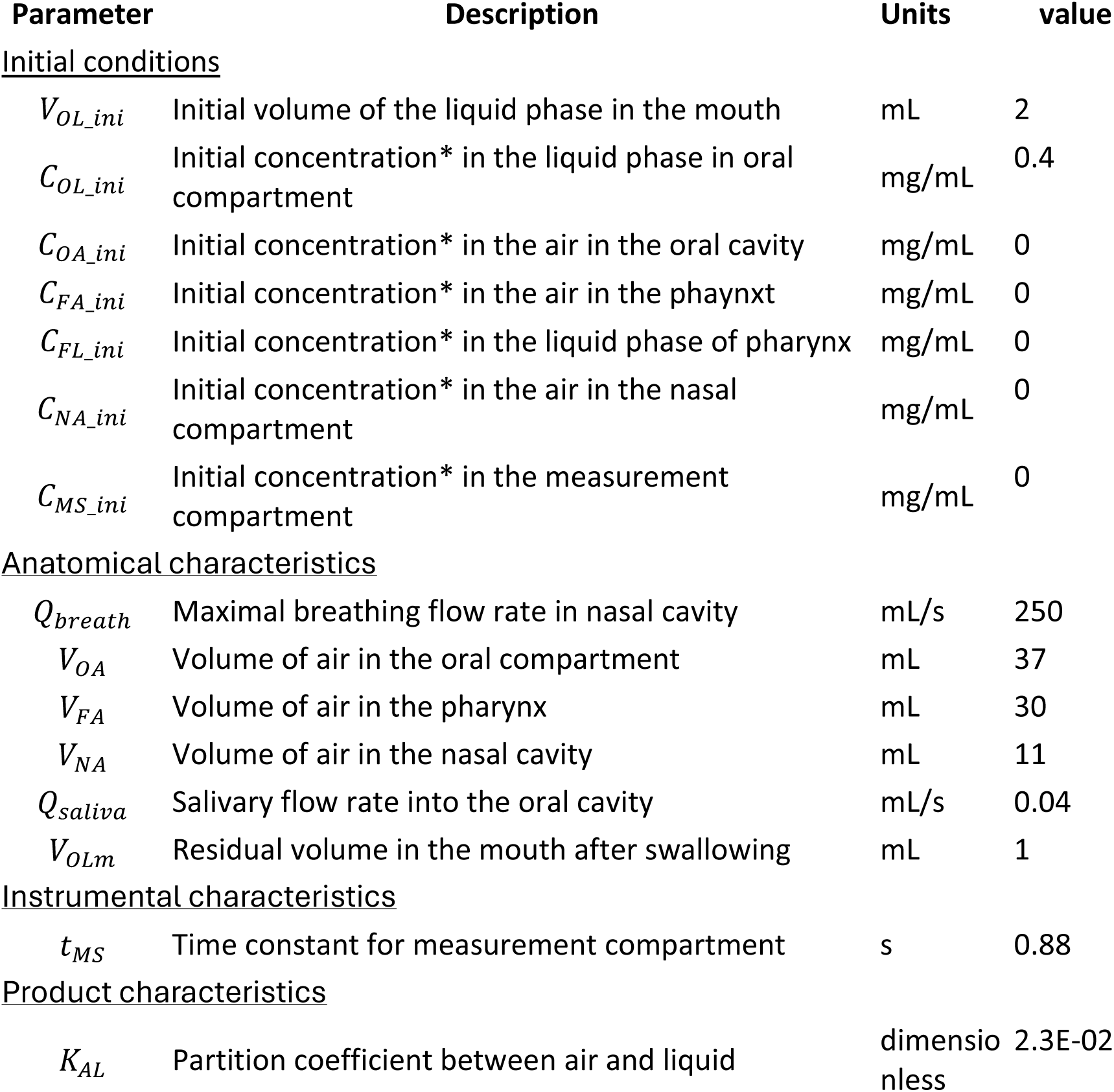
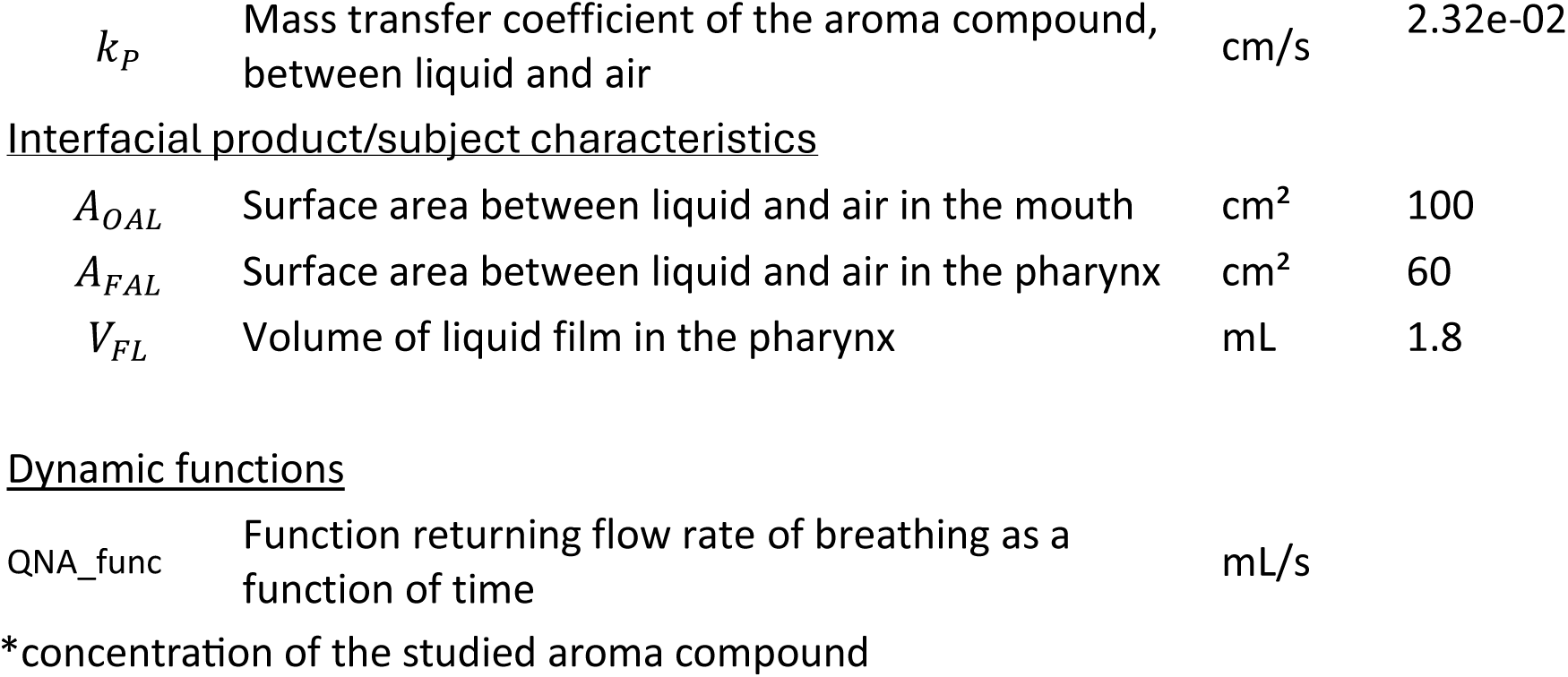
Description of the parameters to be included in the model for tasting an aromatized liquid.

| Parameter | Description | Units | value |
| --- | --- | --- | --- |
| <u>Initial conditions</u> |  |  |  |
| $V_{OL\_ini}$ | Initial volume of the liquid phase in the mouth | mL | 2 |
| $C_{OL\_ini}$ | Initial concentration* in the liquid phase in oral compartment | mg/mL | 0.4 |
| $C_{OA\_ini}$ | Initial concentration* in the air in the oral cavity | mg/mL | 0 |
| $C_{FA\_ini}$ | Initial concentration* in the air in the pharynx | mg/mL | 0 |
| $C_{FL\_ini}$ | Initial concentration* in the liquid phase of pharynx | mg/mL | 0 |
| $C_{NA\_ini}$ | Initial concentration* in the air in the nasal compartment | mg/mL | 0 |
| $C_{MS\_ini}$ | Initial concentration* in the measurement compartment | mg/mL | 0 |
| <u>Anatomical characteristics</u> |  |  |  |
| $Q_{breath}$ | Maximal breathing flow rate in nasal cavity | mL/s | 250 |
| $V_{OA}$ | Volume of air in the oral compartment | mL | 37 |
| $V_{FA}$ | Volume of air in the pharynx | mL | 30 |
| $V_{NA}$ | Volume of air in the nasal cavity | mL | 11 |
| $Q_{saliva}$ | Salivary flow rate into the oral cavity | mL/s | 0.04 |
| $V_{OLm}$ | Residual volume in the mouth after swallowing | mL | 1 |
| <u>Instrumental characteristics</u> |  |  |  |
| $t_{MS}$ | Time constant for measurement compartment | s | 0.88 |
| <u>Product characteristics</u> |  |  |  |
| $K_{AL}$ | Partition coefficient between air and liquid | dimensionless | 2.3E-02 |
| $k_P$ | Mass transfer coefficient of the aroma compound, between liquid and air | cm/s | 2.32e-02 |
| <u>Interfacial product/subject characteristics</u> |  |  |  |
| $A_{OAL}$ | Surface area between liquid and air in the mouth | cm<sup>2</sup> | 100 |
| $A_{FAL}$ | Surface area between liquid and air in the pharynx | cm<sup>2</sup> | 60 |
| $V_{FL}$ | Volume of liquid film in the pharynx | mL | 1.8 |

Dynamic functions
|  |  |  |
| --- | --- | --- |
| QNA_func | Function returning flow rate of breathing as a function of time | mL/s |

\*concentration of the studied aroma compound

#### 2.1.2. Differential Equations

From these initial values, the kinetic of each state variable can be deduced by solving a system of differential equations. Each equation arises from a combination of mass conservation, boundary layer theory, partitioning between phases, and convective transport, reflecting the physiological processes of release, mixing, and advection in the oral and food compartments. The model assumes well-mixed compartments and instantaneous local equilibrium at interfaces, with discrete events (jaw moving and swallowing) handled separately to account for rapid volume and concentration changes. Jaw moving is particularly important because it induces velum opening and release of aromatized air from the mouth to the pharynx and nose (Buettner et al., 2002).

##### 2.2.1.1. Volume of liquid in oral cavity (V_OL_)

The volume of liquid product (or bolus) in oral cavity changes primarily due to salivary secretion.

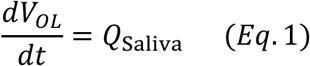

The salivary flow rate, *Q*_Saliva_, adds volume to the bolus in oral compartment. This term represents a simple conservation equation.

##### 2.2.1.2. Oral Compartment (C_OL_)

The concentration of volatile compounds in the oral liquid decreases due to two mechanisms: (i) transfer to oral air, symmetric to the gain term in *C_OA_*, ensuring mass conservation and (ii) dilution by saliva, proportional to salivary flow rate *Q_Saliva_*.

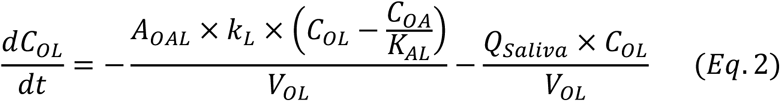

This reflects the physical reality that saliva continuously dilutes the food bolus and reduces volatile availability.

##### 2.2.1.3. Oral Air Compartment (C_OA_)

The oral cavity contains an air phase in which volatile compounds are released from the food bolus. The driving mechanism is the deviation from thermodynamic equilibrium between the liquid phase located in the oral cavity (*C_OL_*) and the oral air phase (*C_OA_*).

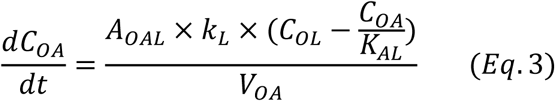

##### 2.2.1.4. Pharyngeal Air Compartment (C_FA_)

The pharyngeal air compartment is a key junction between oral release and nasal perception. The first term models volatile release from the product located in the pharynx (*C_FL_*) into the pharyngeal air. The second term represents airflow-driven exchange between the pharynx and the nasal cavity, controlled by *Q_NA_*, due to breathing. The piecewise formulation accounts for bidirectional airflow: positive *Q_NA_*: transport from nose to pharynx; negative *Q_NA_*: retronasal transport from pharynx to nose.

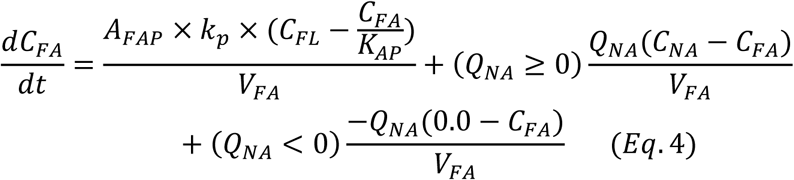

##### 2.2.1.5. Pharyngeal Product Compartment (C_FL_)

The liquid bolus present in the pharynx releases volatile compounds into the surrounding air.

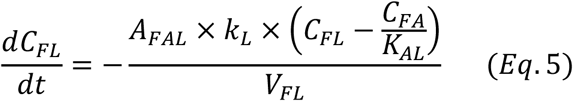

##### 2.2.1.6. Nasal Compartment (C_NA_)

The nasal cavity is modeled as a non-absorbing air compartment whose concentration changes are driven exclusively by airflow.

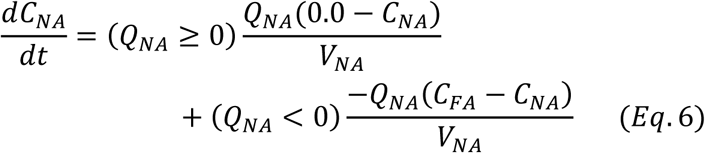

During inspiration (*Q_NA_* ≥ 0), clean air enters and dilutes nasal volatiles. During expiration or retronasal transport (*Q_NA_* < 0), volatile-rich air from the pharynx enters the nasal cavity. This formulation captures the directionality of nasal airflow during breathing.

##### 2.2.1.7. PTR-MS Compartment (C_MS_)

This equation is equivalent to a first-order linear transfer process from the nasal cavity to the measuring instrument.

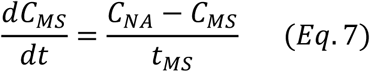

#### 2.2.2. Events

In addition to continuous transport and mass transfer processes, the model incorporates discrete physiological events that induce rapid redistribution of volatile compounds between compartments. These events are implemented as instantaneous state updates and account for mechanical mixing and bolus displacement occurring during eating. Two types of events are considered: jaw movement and swallowing.

##### 2.2.2.1. Jaw movement

The jaw movement event represents the mechanical mixing of the air in the three gas compartments (oral, pharyngeal and nasal compartments).

A volume-weighted average concentration is first computed for homogenizing the concentration in these compartments:

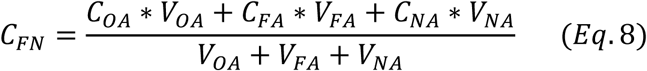

The system state is then instantaneously updated as follows: *C_OA_*←*C_FN_*; *C_FA_*←*C_FN_*; *C_NA_*←*C_FN_*. All other state variables remain unchanged.

##### 2.2.2.2. Swallowing

The swallowing event represents the clearance of the oral cavity and the transfer of the bolus toward the pharynx and esophagus. In practical terms, the volume of the food bolus in the oral cavity is reduced to a minimal value (*V_OLm_*), its concentration is transferred to the pharyngeal compartment, and the air-phase concentration is homogenized across the different air compartments.

The system state is then instantaneously updated as follows: *C_OA_*←*C_FN_*; *C_FA_*←*C_FN_*; *C_NA_*←*C_FN_*; *C_FL_*←*C_OL_*, *V_OL_*←*V_OLm_*where *C_FN_* is given by eq. 8.

### 2.3. Modelling aroma release from a solid product

A separate model was developed for solid foods. It is conceptually similar to the solution model, but includes additional compartments and processes specific to solid matrices.

- An additional solid product is represented in the mouth
- This solid is dissolved into saliva. This step is controlled by the contact surface area between the product and saliva, and the rate of dissolution of the solid product
- The contact surface area is modelled as depending on the volume of the solid product and on the shape parameter of the solid product (depending on potential chewing)
- The speed of dissolution of the product varies according to its temperature
- The temperature of the solid is varying proportionally to the difference of temperature between the product and the mouth
- The rest of the phenomena are similar to the solution case (aroma release of solution in the air in the oral cavity, then in the pharynx, nasal cavity and finally to PTR-MS)
- Moreover, swallowing may occur either with complete ingestion (“total_swallow”) of the solid product or without swallowing it (“swallow”). In this case, only the saliva is swallowed. Both events are implemented in the model.

#### 2.3.1. Parameter and variable description

Considering solid foods leads to an additional solid compartment for the product whose volume is *V_OP_* (considered as a state variable, decreasing along time due to melting). The liquid compartment volume is *V_OL_* and is constituted of the saliva and the melted part of the solid food, with varying aroma concentration *C_OL_*. The temperature of the product *T_P_* and the shape parameter of the product *S_P_* (developed below) are additional state variables. The other state variables are similar to the solution case *C_OA_*, *C_FA_, C_NA_, C_MS_.* They depend on the parameters described in Figure 2 and Table 2.

**Figure 2:**
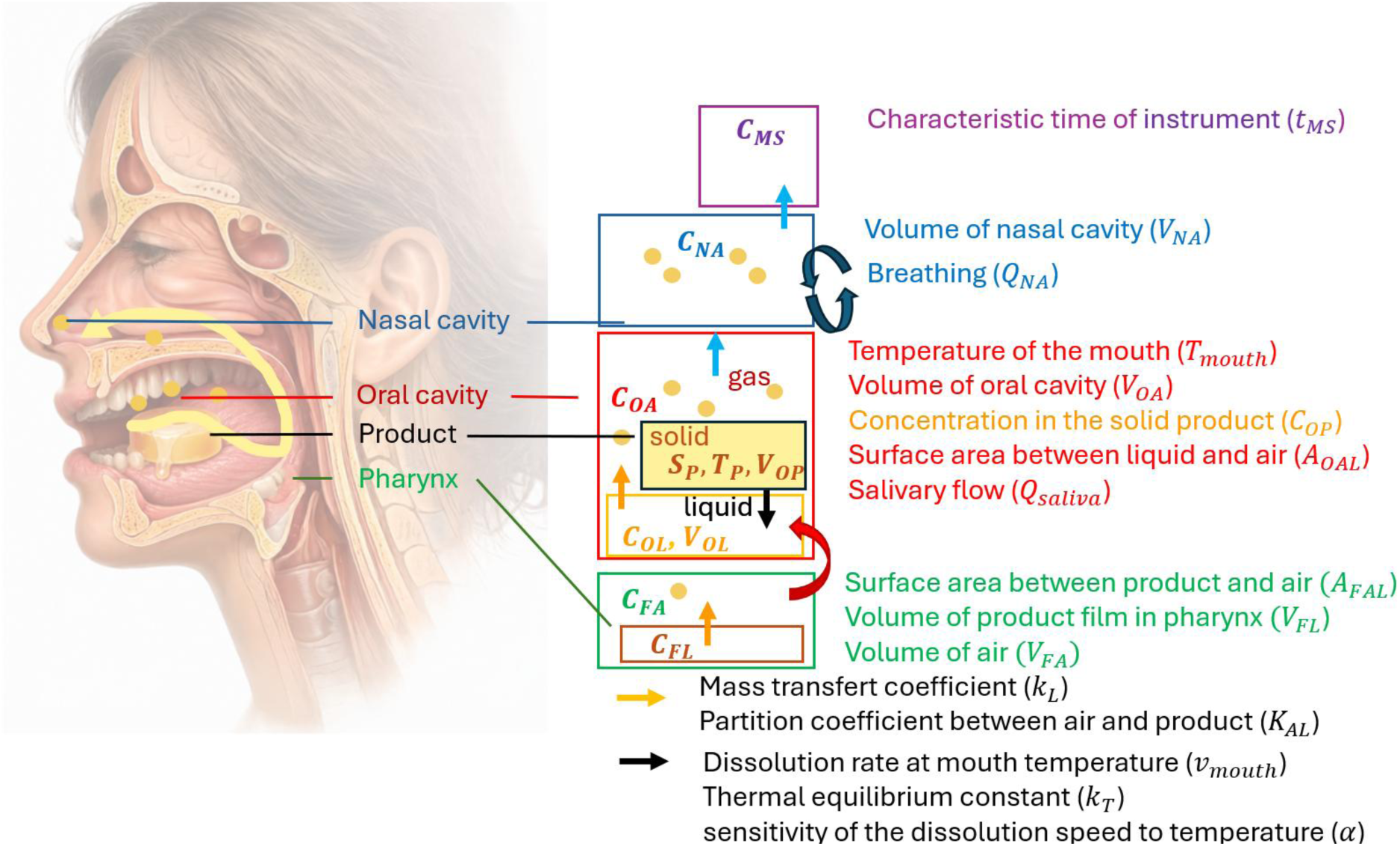
Representation of the modeling of aroma release from a melting product in the mouth and related variables and parameters.

**Table 2:** list of the parameter’s values for modeling aroma release from a solid product.

| parameter | description | unit | value |
| --- | --- | --- | --- |
| <u>Initial conditions</u> |  |  |  |
| $V_{OP\_ini}$ | Initial volume of the solid product in the mouth | mL | 2 |
| $C_{OL\_ini}$ | Initial concentration of the aroma in the liquid phase in the oral compartment | mg/mL | 0 |
| $C_{OA\_ini}$ | Initial concentration of aroma in the air in the oral cavity | mg/mL | 0 |
| $C_{FA\_ini}$ | Initial concentration of aroma in the air compartment of the pharynx | mg/mL | 0 |
| $C_{FL\_ini}$ | Initial concentration in the liquid phase of pharynx | mg/mL | 0 |
| $C_{NA\_ini}$ | Initial concentration of aroma the air in nasal cavity | mg/mL | 0 |
| $C_{MS\_ini}$ | Initial concentration in of aroma the measurement compartment | mg/mL | 0 |
| $T_P\_ini$ | Initial temperature of the product | °C | 4 |
| $S_P\_ini$ | Initial shape parameter | - | 1.27 |
| <u>Anatomical characteristics</u> |  |  |  |
| $V_{OA}$ | Volume of oral compartment | mL | 37 |
| $Q_{breath}$ | Maximal breathing flow rate in nasal cavity | mL/s | 250 |
| $Q_{saliva}$ | Salivary flow rate into oral cavity | mL/s | 0.04 |
| $V_{FA}$ | Volume of air in pharynx | mL | 30 |
| $V_{NA}$ | Volume of air in the nasal cavity | mL | 11 |
| $T_{mouth}$ | Temperature of the mouth | °C | 36 |
| <u>Instrumental characteristics</u> |  |  |  |
| $t_{MS}$ | Time constant for the instrument response | s | 0.88 |
| <u>Product characteristics</u> |  |  |  |
| $K_{AL}$ | Partition coefficient between air and liquid phase (saliva with dissolved product) | dimensionless | 2.3E-02 |
| $C_{OP}$ | Concentration of aroma in the solid product | mg/mL | 0.4 |
| $k_L$ | Mass transfer coefficient between solution and air for the aroma | $cm\ s^{-1}$ | 2.32e-02 |
| $V_{FL}$ | Volume of liquid film in pharynx | mL | 1.8 |
| $v_{mouth}$ | Speed of dissolution of the product at the temperature $T_{mouth}$ | cm/s | 0.015 |
| $\alpha$ | sensitivity of the dissolution velocity to temperature | dimensionless | 0.5 |
| $k_T$ | thermal equilibration constant of the product in the mouth | $s^{-1}$ | 0.1 |
| <u>Interfacial product/subject characteristics</u> |  |  |  |
| $A_{OAL}$ | Surface area between liquid and air in the mouth | cm<sup>2</sup> | 100 |
| $A_{FAL}$ | Surface area between liquid and air in pharynx | cm<sup>2</sup> | 60 |
| $\lambda_{chew}$ | Factor for chewing constant | - | 1.25 |
| <u>Dynamic functions</u> |  |  |  |
| Qna_func | Function returning flow rate breathing | mL/s |  |

#### 2.3.2. Dynamic equations

##### 2.3.2.1. Shape factor

*S_P_* is a dimensionless shape factor describing the geometry and surface development of the product. For an ideal homogeneous dissolution without structural changes the shape factor remains constant. However, oral processing modifies the product structure through successive mastication events. Although chewing does not directly change the remaining product volume, it increases the available contact surface by fragmenting the product into smaller pieces and creating new interfaces with saliva. To account for these structural changes, the shape factor is introduced as an additional state variable. Between mastication events, it is assumed to remain constant:

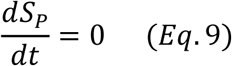

*A_OLP_* is the contact surface area between the solid product and the oral liquid. *A_OLP_* is assumed to depend on the product volume according to

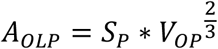

Chewing event excepted, the relationship between volume and area of the product assumes isotropic dissolution and a constant solid shape throughout the process. Note that the initial value of *S_P_*_,*ini*_ is :

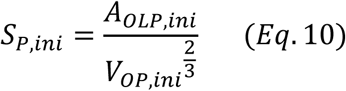

##### 2.3.2.2. Solid product dissolution (V_OP_)

The solid product dissolves at the solid–liquid interface. The rate of solid volume decrease is assumed to be proportional to the contact area between the solid product and the oral liquid and to a temperature-dependent surface recession velocity:

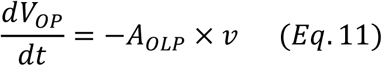

##### 2.3.2.3. Temperature

The surface recession velocity is assumed to depend linearly on temperature according to:

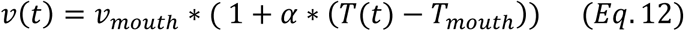

where *v_mouth_* (*cm*. *s*^−1^) is the surface recession velocity of the solid product at the mouth temperature *T_mouth_*, and *α* (°C^−1^) quantifies the sensitivity of the dissolution velocity to temperature.

The product temperature is assumed to relax exponentially toward the mouth temperature:

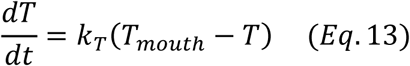

where *k_T_* is the thermal equilibration constant (in *s*^−1^).

##### 2.3.2.4. Saliva Volume Dynamics (V_OL_)

The oral liquid volume consists of saliva and dissolved product. The oral liquid volume increases due to both saliva secretion and solid dissolution:

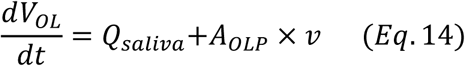

The dissolved product volume is assumed to increase by the same amount as the solid product volume decreases, assuming complete conversion of the solid into dissolved material.

##### 2.3.2.5. Concentration in Oral Liquid Phase (C_OL_)

This equation is a mass balance on the aroma compound in the mouth. It accounts for (i) input from solid dissolution, (ii) loss to the oral air phase via mass transfer, (iii) dilution due to changing liquid volume.

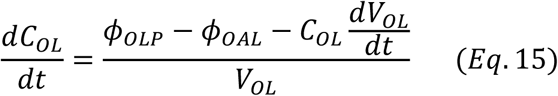

where:

- *φ_OLP_* = *A_OLP_* × *v* × *C_OP_* (solid → liquid aroma flow): assuming a uniform product flavoring.
- 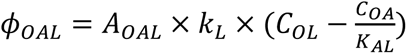 (liquid→gas aroma flow) where *A_OAL_* is the oral liquid–air interfacial area, *k_OL_* is the mass transfer coefficient and *K_AL_* is the air/liquid partition coefficient

The equations used for the remaining state variables *C_OA_*, *C_FA_*, *C_FL_*, *C_NA_*, *C_MS_* are identical to those presented in 2.2 for the liquid consumption model.

#### 2.3.3. Event Functions (Discrete State Updates)

##### 2.3.3.1. Chew event

This event represents the opening of the velopharynx whose main effect is in this case a fast air flow and an instantaneous mixing of the air in the mouth, pharynx and nose as the liquid model (*2.2.2.1)*. This event induces air mixing between various compartments without loss of aroma.

Furthermore, the shape parameter increases by being multiplied by *λ_c_*_ℎ*ew*_(>1)

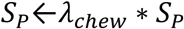

The other state variables of the model remain unchanged.

##### 2.3.3.2. Partial swallowing event

This event represents a partial swallowing of the liquid phase in the mouth, where oral and pharyngeal air compartments rapidly mix but the solid product is not swallowed. It consists in a ‘chew’ event with the following additional actions:

*V_OL_*←*V_OLm_*; *C_FL_*←*C_OL_* where *V_OLm_* is the part of liquid that remains in the mouth after swallowing.

##### 2.3.3.3. total_swallow event

total_swallow represents a ‘swallow step’ with an additional action: the solid product is entirely swallowed: *V_OP_*←0.

### 2.4. Simulations

#### 2.4.1. *In vivo* simulations for liquid foods

The parameter values used in the simulations are presented in the “value” column of Table 1 and were chosen to match those used in the experimental setup described later.

Breathing was simulated as a sinusoid *Q_NA_*(*t*) = *Q_Breath_* ∗ sin (2*π*(*t* + 5)/5) where *Q_Breath_* = 250 mL/s. Two events of chewing at 20 and 40 s, then two swallowing events at 60 and 80 were implemented.

#### 2.4.2. *In vivo* simulation for solid foods

The parameter values used in the simulations are presented in the “sim” column of Table 2 and were chosen to match those used in the experimental setup described later. Similarly to the solution case, breathing was simulated with the same sinusoid as 2.4.1.

One event of ‘chew’ at 40 s, then one partial swallowing event at 60 and a total swallowing at 80 s were implemented.

### 2.5. Comparison with experimental data

#### 2.5.1. Presentation of the data

Experimental data was collected to compare the results of these models to reality. *In vitro* data was presented in Andriot *et al*. (2026a) and *in vivo* data was documented in (Andriot *et al*., 2026b).

Thirty subjects participated in a sensory study during which they consumed a product over a two-minute period while following standardized oral processing instructions and being connected to PTR-MS, an instrumental device that allows aroma intensity (proportional to concentration) in the nose to be quantified with high temporal resolution (0.1 s). Participants were instructed to report each action (jaw movement, chew, saliva swallow or product swallow) by pressing a designated key on a computer keyboard. Two products were evaluated, each tested six times per subject: a liquid solution and a semi-solid gelatin gummy disc, both formulated with isoamyl acetate at comparable concentrations (0.4 g·L⁻¹).

Each tasting session was divided into three successive phases, depending on the product. For the solution, participants first remained inactive for 30 s, followed by a 30 s phase during which they moved their jaws and reported each biting action. Finally, the entire liquid was swallowed, after which participants performed either frequent swallowing (‘freq’ protocol, three repetitions) or infrequent swallowing (‘rare’ protocol, three repetitions). For the gelatin gummy discs, participants first remained inactive for 30 s, then either masticated (‘chew’ protocol, three repetitions) or sucked (‘succ’ protocol, three repetitions) the product, before swallowing and subsequently swallowing at a regular rate (Andriot et al., 2026b). The different food oral processing (FOP) (‘freq’ and ‘rare’ for the solution, ‘succ’ and ‘chew’ for the gel) were compared by plotting the average release curves.

#### 2.5.2. Simulations in the same conditions as collected experimental data

##### 2.5.2.1. Simulation of solution tasting in experimental conditions

We used here the same parameters for all subjects, presented in the ‘value’ column of table 1. Product parameters were derived from the experiment (concentration, volume), as well as from previous estimations (transfer coefficients, see Peltier et al., 2027)) or additional experiments using GC–MS (partition coefficients).

The breathing signal was estimated using the isoprene ion, which served as a marker of respiration. High isoprene concentrations measured in the nose correspond to expiration, whereas values close to zero correspond to inspiration. However, in the code, the nasal airflow convention is reversed, with positive values indicating inspiration and negative values indicating expiration. Therefore, the estimated breathing signal was first centered and then sign-inverted to align it with the nasal airflow convention. The resulting signal was then normalized by its maximum absolute value to obtain values ranging from −1 to 1 and finally scaled by a representative breathing flow reported in the literature (*Q_Breath_*=250 mL/s).

Each subject reported the timing of events (‘*jaw_move*, ‘*swallow*’). The reported timestamps for *jaw_move* were used directly. In contrast, because swallowing physiologically occurs during the expiratory phase and is typically followed by an expiration (so-called ‘swallow breath’), the reported swallowing times were adjusted to coincide with the onset of the subsequent expiration.

We first visualized the results for one given subject (simulated and experimental curves of aroma release, breathing signal). To visualize and assess the results for all evaluations (90 curves), we compute the experimental and modelled curve average (at panel level) after min-max standardization.

We then calculated the RMSE and the corresponding R² between the averaged and normalized experimental and modelled curves. For this purpose, each of the two observed average curves (on replicate and subjects for one given protocol) was normalized by the maximum value observed across the two curves on 30-120 s period. This normalization was necessary because of the multiplicative relationship between the actual aroma concentration and the number of counts measured by PTR-MS is not known.

##### 2.5.2.2. Simulation of the gel data with experimental settings

We used here the same parameters for all subjects, presented in the ‘value’ column of table 2. The breathing estimation and swallowing adjustments were computed similarly to 2.5.3.1. In the chewing protocol, the dissolution rate (*v_mouth_*) was chosen as 0.15 mm/s. The value of *S_P_*__*ini*_ (=1.27) was defined as the ratio of the initial solid–liquid interfacial area (here, πr², where r is the radius of the gelatin gummy disc, 8mm) to the initial product volume (2 mL) raised to the power of 2/3. RMSE and R2 were also calculated similarly to 2.5.3.1.

We first visualized the results for one given subject (simulated and experimental curves of aroma release, breathing signal). Then, we computed the experimental and modeled curve average (at panel level) after min-max standardization for all experiments (90 curves).

### 2.6. Computational aspects

All the models presented in this paper were implemented in digimouth (existing_models.py) presented in Peltier (2027). The code required for reproducing the figures of the paper is also available in digimouth package. All calculations were computed with Python 3.12.1.

## 3. Results

### 3.1. Simulations

#### 3.1.1 *In vivo* simulation for liquid solution intake

The example simulation for solutions is presented in Fig 3.

**Figure 3:**
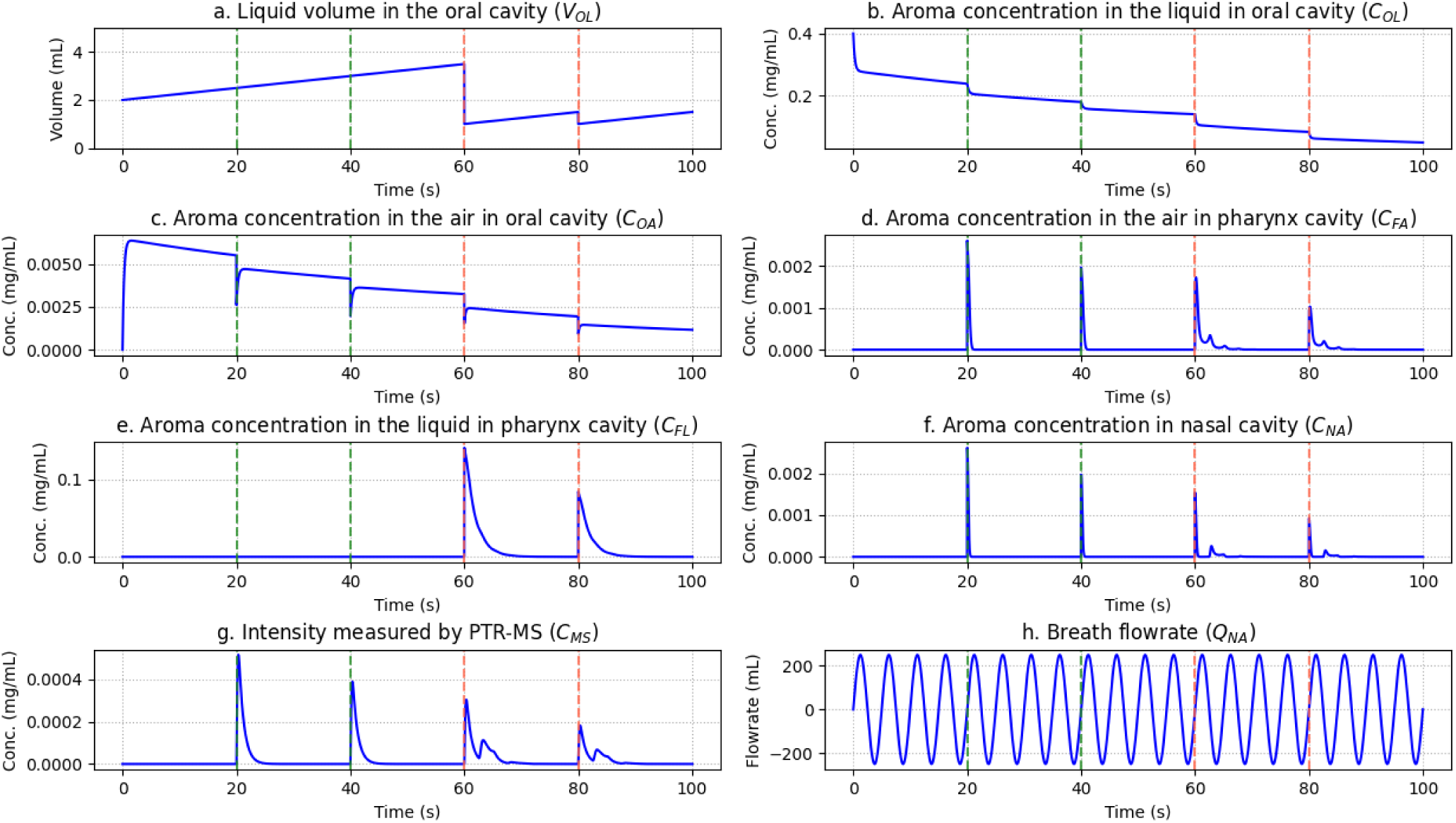
simulation of liquid with four events: jaw moves (green) at 20 s and 40 s and swallows at 60 and 80 s (red) in the different compartments.

First, the volume of the bolus (product + saliva) increases due to the constant production of saliva (Fig 3a). After the first swallowing event (60 s), only a small fraction of the product (*V_OLm_*) remains in the mouth; salivary flow then causes the volume to increase again, and the second swallowing event (80 s) reduces it back to *V_OLm_*. Regarding the aroma concentration in the liquid, it decreases over time as aroma is released in the oral cavity and the liquid diluted by fresh saliva (Fig 3b). The aroma concentration in the air in the oral cavity increases due to release from the liquid, reaching a maximum at around 2 seconds, and then slowly decreases, following the concentration in the liquid, up to the partition factor (Fig 3c). After pseudo-equilibrium between air and liquid in the oral cavity is reached, aroma dilution by saliva becomes the dominant process, leading to a gradual decrease in concentration driven mainly by saliva inflow in the oral cavity. At chewing time (t=20 s and t=40 s) the gas is mixed among the three cavities (oral, pharyngeal, and nasal, Fig 3d). A similar phenomenon occurs at t=60 s and 80s, during swallowing events. The aroma concentration in the pharynx liquid layer remains zero until the first swallow, after which it is reset to the oral concentration and quickly depleted, due to losses to the pharynx air (Fig 3e). The concentration in the nasal cavity essentially follows that of the pharyngeal air (Fig 3f). Finally, the PTR-MS signal exhibits broader peaks due to instrumental response dynamics (low-pass filtering, Fig 3g). Finally, recall that the simulation was performed assuming a sinusoidal breath flow rate (Fig 3h).

#### 3.1.2 *In vivo* simulation for solid product intake

Three events occur during this simulation: a ‘chew’ at 40 s, a swallow at 60 s and a total swallow at 80 s. Figure 4 illustrates the simulation for solid product intake.

**Figure 4:**
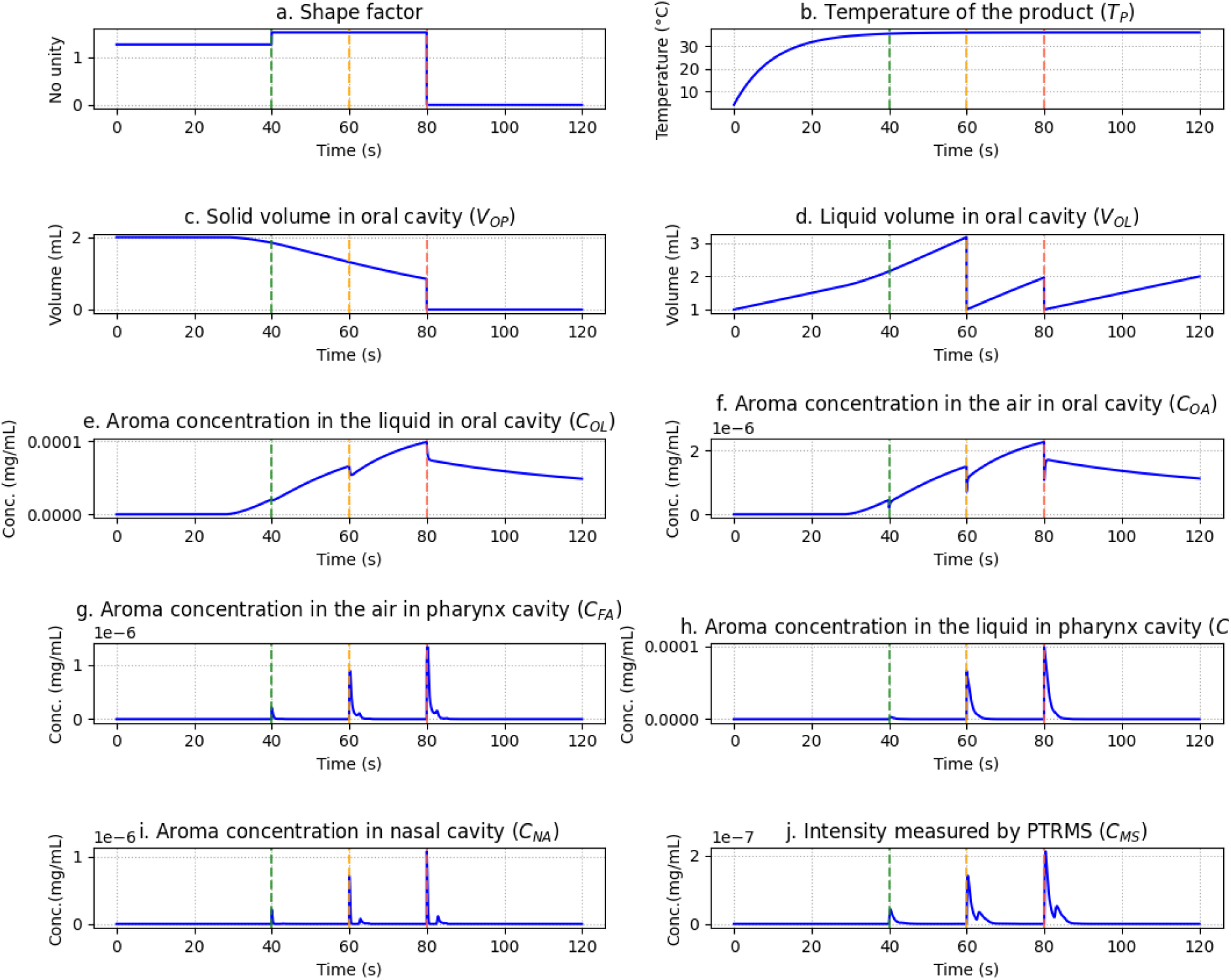
Simulation results for solid product in the different compartments. Green dotted lines indicate chewing events, orange dotted lines indicate partial swallowing events, and red dotted lines indicate complete swallowing events.

First, the shape factor is multiplied by 1.25 at the ‘chew’ time, then is constant and finally set to zero when the solid product is totally swallowed (Fig 4a). The temperature of the product increased from its initial storage value to the mouth temperature (Fig 4b). The volume of oral solid product decreased due to its melting at a gradually increasing rate (Fig 4c), due to product temperature increase. At 80 seconds the product is totally swallowed and disappears, as expected. The volume of liquid constantly rises due to the constant production of salivary flow, then falls to a minimal volume at each swallowing (60 and 80 s, Fig 4d). After each swallowing event, only a small fraction of the liquid product remains in the mouth *V_OLm_* = 1 *mL*; salivary flow then causes the volume to increase again. Regarding the aroma concentration in the liquid phase, it increases in parallel with product (gummy disk) dissolution and sharply decreases at each swallow due fast release to the air (Fig 4e). After the total swallow (80 s) it slowly decreases due to dilution with fresh saliva. The aroma concentration in the air in the oral cavity follows that in the liquid, due to quick transfer dynamics, as for solution (Fig 4f). Then, when chewing (t=40 s), the gas is distributed among the three cavities (oral, pharyngeal, and nasal, Fig 4g). A similar phenomenon occurs during swallowing events at t=60 s and t=80 s. For the pharynx, nose and PTR-MS compartments, the trends are similar to the liquid solution (Fig 4g-j).

### 3.2. Comparison with experimental data

#### 3.2.1. Solution in experimental conditions

Figure 5 presents the individual experimental data and event annotations (a), the corresponding simulated data (b), together with the breathing signal (c) for a selected panelist. Graphs (d) and (e) show the mean curves with their standard errors for the experimental and simulated data, respectively.

**Figure 5:**
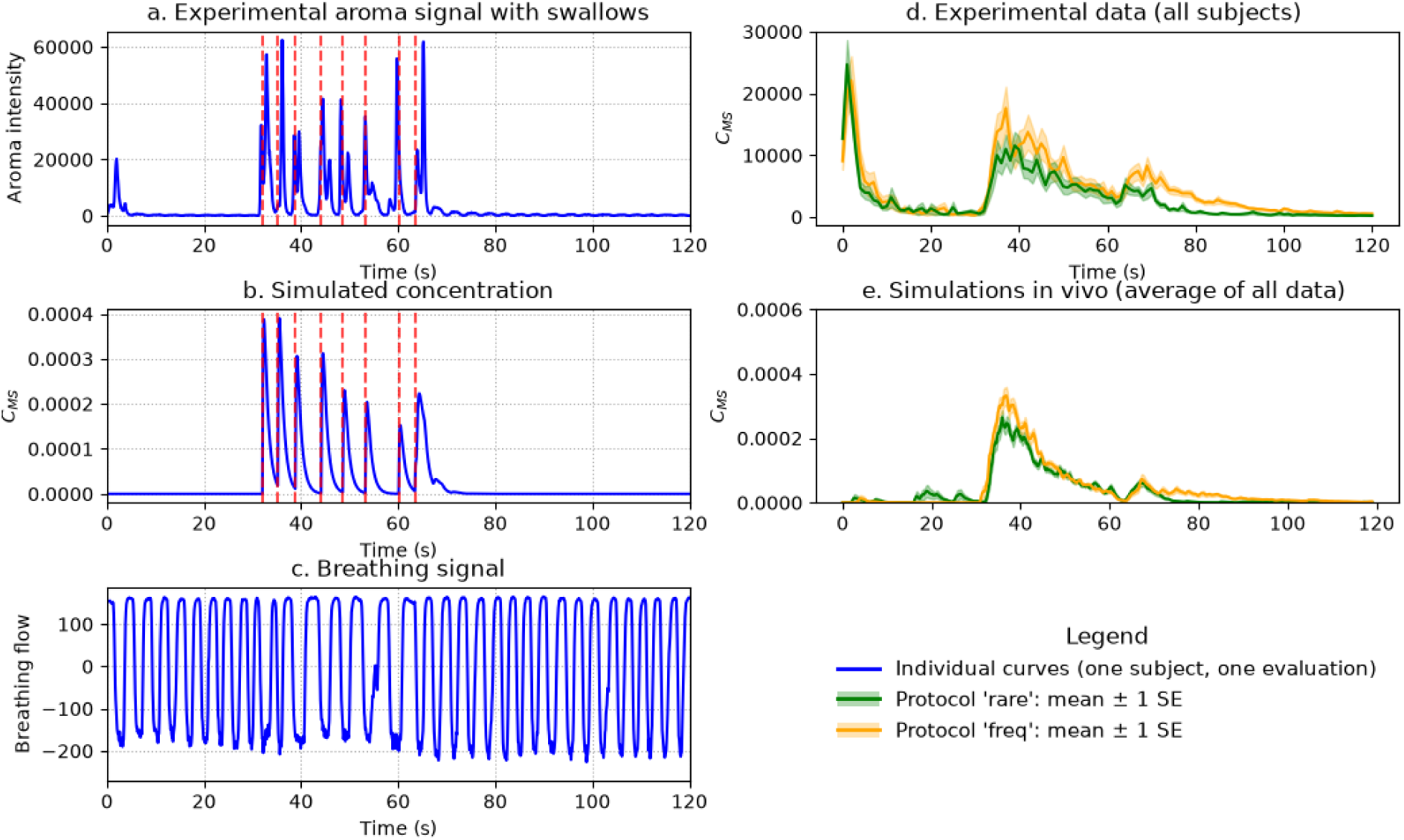
Example of experimental (a) and corresponding simulated aroma release (b) and experimental breath signal (c) for one replicate of one individual. Averaged and standard error of curves for experimental data (d) and simulated data (e) for the protocol with few swallows after 60 s (‘rare’, green line) and with frequent swallows after 60 s (‘freq’, orange line)

Overall, the simulations reproduce the main features of the experimental curves. The differences between the two protocols (with low or high number of swallows after 60 s) are similar in experimental and simulated data.

The ratio between the aroma peak around 35 s and the peak around 65 s is slightly different between the protocols in the experimental data and well reproduced in simulation.

One notable difference is the presence of a peak around 5 s in the experimental data, which is absent from the simulations. This peak most likely corresponds to an orthonasal response occurring when participants brought the sample close to their nose before placing it in the mouth. As this phenomenon was not included in the model, it is not reproduced in the simulations.

Another difference is that the simulated signal returns close to zero shortly after 60 s following swallowing, whereas the experimental signal decreases more gradually. This discrepancy is likely due to the swallowing event being represented as instantaneous and perfectly synchronized in the simulations, while swallowing occurred over a finite duration and with greater variability among participants.

After normalizing both experimental and simulated data to a [0, 1] range, the ‘freq’ protocol yielded an RMSE of 0.13 and an R² of 0.70, whereas the ‘rare’ protocol yielded an RMSE of 0.07 and an R² of 0.86. Overall, the model showed a better agreement with the experimental data for the ‘rare’ protocol than for the ‘freq’ protocol, as indicated by the lower RMSE (0.07 vs. 0.13) and higher R² (0.86 vs. 0.70). This suggests that the model more accurately captured the temporal profile of aroma release under the ‘rare’ protocol, which does not include swallowing. The lower agreement observed for the ‘freq’ protocol suggests that the swallowing model could be improved by developing new models of swallowing or a more accurate event reporting.

#### 3.2.2. Gelatin gummy disc in experimental conditions

Simulated and experimental results show similar kinetics, although not identical (Fig 6). For the ‘chew’ protocol, a low level of aroma release is observed before 30 seconds, followed by the onset of release when the subject starts chewing (30–60 s). For the ‘succ’ protocol, the release remains much lower, although nonzero, between 30 and 60 s. After complete swallowing (t = 60 s), a residual aroma release persists with each subsequent swallow, following a roughly exponential decay.

**Figure 6:**
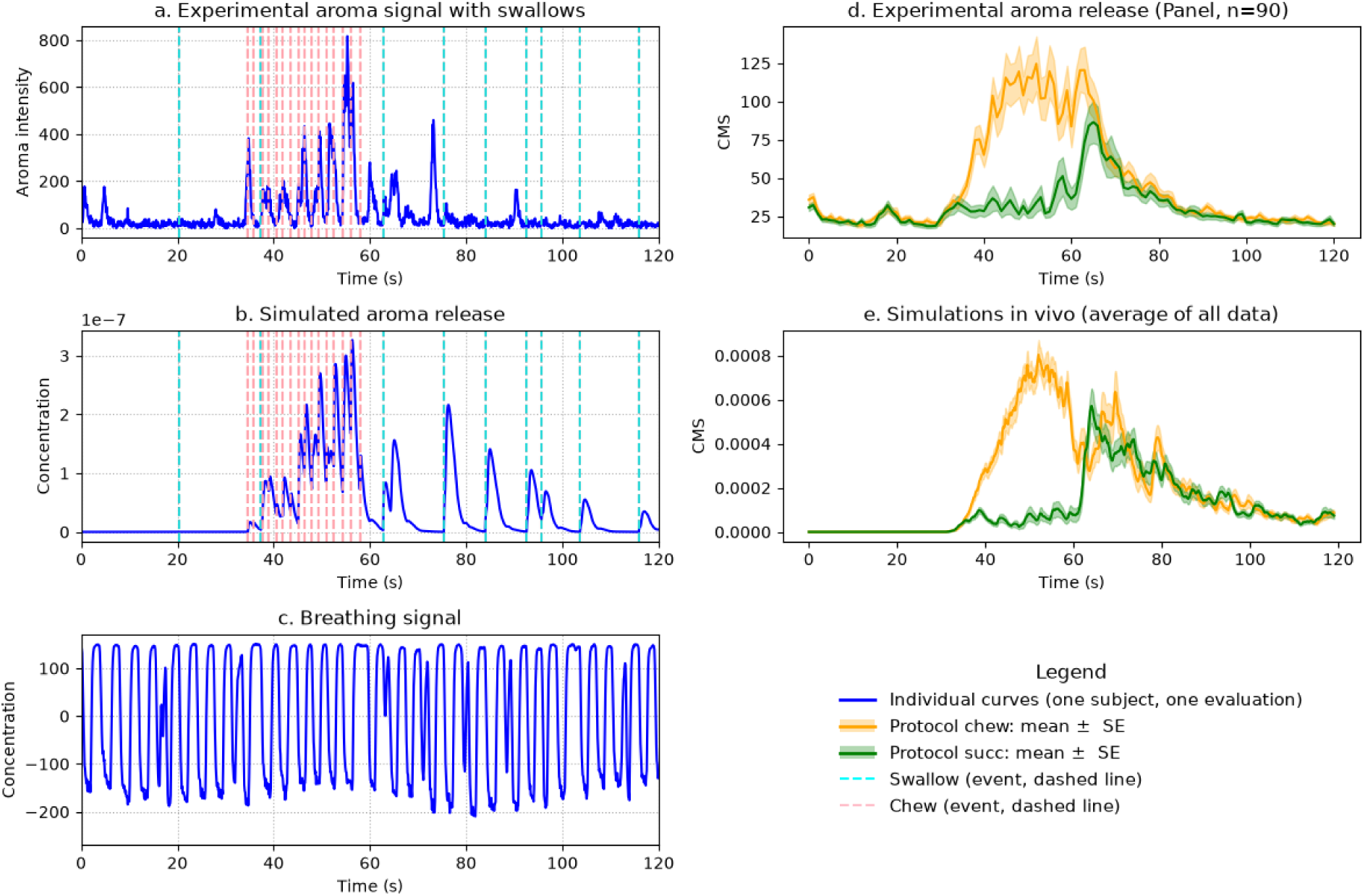
Example of experimental (a) and corresponding simulated aroma release (b) and experimental breath signal (c) for one replicate of one individual, for the ‘chew’ protocol. Averaged and standard error of curves for experimental data (d) and simulated data (e) for the ‘chew’ (in orange) and ‘succ’ (green) protocols.

Comparison of the panel-averaged curves revealed similar overall kinetics for the two protocol conditions. When the panelists were instructed to suck the product (green), aroma release remained relatively low during the 30–60 s period, followed by a sharp peak at (t = 60 s) before gradually returning toward zero. In contrast, when the panelists were instructed to chew the product (orange), aroma release increased after 30 s and then progressively decreased. Overall, these characteristic features were well reproduced by the model.

A slightly different kinetic profile among protocols is observed in simulation during the final stage of chewing (60-90 s). The simulations show two chewing-related peaks that are absent from the experimental data, suggesting that the models could be further refined to better capture the chewing of solid products.

After normalizing both experimental and simulated data to a [0, 1] range, the ‘chew’ protocol yielded an RMSE of 0.18 and an R² of 0.57, whereas the ‘succ’ protocol yielded an RMSE of 0.13 and an R² of 0.31.

## 4. Discussion

### 4.1. Limits of the current models and their related experimental data

Chewing and swallowing are implemented as punctual events assumed to occur at precise moments. However, in practice, these timestamps are often imprecise. For example, in our dataset, swallowing times are self-reported by the panelists, which may introduce uncertainty, delays, or reporting errors. This source of variability is not currently accounted for in the model; however, future improvements such as automatic analysis of video sequences could incorporate methods to address temporal uncertainty in event timing and reduce cognitive load of the panelists, allowing them to concentrate on aroma perception. A more robust approach would be to measure these events automatically using video-based systems, for example to detect jaw movements associated with chewing. Swallowing events are more challenging to capture, but could potentially be identified using physiological signals such as electromyography (EMG), for instance with sensors placed near the laryngeal or glottal region. Such measurements would provide more accurate and objective timing information, but would also require more advanced signal processing, including careful synchronization between multiple devices.

The estimation of the interfacial area between the solid product and the liquid in the mouth is based on a simple formula that may be overly simplistic. Experimental studies could therefore be conducted to better characterize its variation.

From a theoretical standpoint, the rate of thermal equilibration between the solid product and the mouth, accounted in the model by the coefficient *k_T_*, should be proportional to the contact area between the oral liquid and the product *A_OLP_* and inversely proportional to the solid product volume (*V_OP_*). A more rigorous formulation of the heat balance would therefore be:

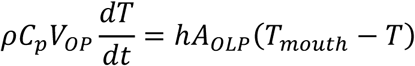

where *ρ*, *C_p_*, and ℎ are assumed to remain approximately constant. This formulation implies that the effective temperature equilibration coefficient scales with the ratio *A_OLP_*/*V_OP_*. However, introducing this dependency is not expected to substantially affect the simulation results under the conditions investigated in this study, while it may lead to numerical instabilities as *V_OP_* reaches zero. For these reasons, the simpler formulation was retained.

In this study, the data were normalized to focus solely on the kinetics. However, the amplitude of the signal is also of interest. The relationship between the actual concentration and the signal recorded by the PTR-MS device is not straightforward. Further work could aim to quantify this relation through experimental investigations.

Breathing was quantified using isoprene as a marker, which is useful for identifying inspiration and expiration events. However, the signal intensity reflects individual metabolic differences rather than the airflow, as some subjects naturally produce higher amounts of isoprene than others. Future work could explore alternative approaches for characterizing the respiratory signal, such as combining acetone and isoprene measurements or employing other types of respiratory monitoring techniques such as breathing move detection.

### 4.2. Discussing experimental results

#### 4.2.1. Comparing experimental and simulated results

Regarding the solution, the occurrence of an orthonasal peak could also be considered in the model. This could be accounted for by introducing a “sniffing” event; however, this event would depend on several uncontrolled factors that were not measured in this study, such as the distance between the cup and the nose, or the exact timing of sniffing. A more appropriate correction would have been to instruct participants not to smell the sample, or alternatively to administer the solution using a straw to prevent the occurrence of an orthonasal peak.

Regarding the gelatin gummy disc, the small dip in the simulated aroma release observed around 60 s for chewing protocol could coincide with a lower number of chewing events reported by the panelists immediately before swallowing. In this case, one possible explanation is that participants may have focused on the upcoming swallowing event and consequently under-reported chewing events. This highlights a limitation of relying on manually reported oral events and further supports the interest of objective, automated detection methods for chewing and swallowing.

#### 4.2.2. Event times

In this study, event timings were self-reported by participants, which introduces several sources of bias, including temporal misalignment (delays or anticipation), and missing or inaccurate declarations.

### 4.3. Improving the models

#### 4.3.1. Choice of parameters and model

Some parameter choices were made in an admittedly arbitrary manner and could be further optimized or directly measured experimentally. Similarly, the modeling assumptions adopted here are open to discussion. The software package allows users to modify these assumptions and test alternative parameter sets and model structures, in order to improve both parameter estimation and model fidelity, and ultimately advance understanding in the field of aroma release.

#### 4.3.2. Choice of events

During mastication, the exchange surface between the solid food and the liquid phase increases. This effect is accounted by multiplying the area by a constant factor at each bite. However, an alternative model can be implemented that allows the user to define the kinetics of the exchange surface evolution, and to relate it to chewing events in a more elaborate way.

It is also possible for users of digimouth to add their own events (opening the mouth, moving the product…)

#### 4.3.3. Adding subject heterogeneity

The only simulation inputs that varied between subjects in this study were the reported timing of events and the breathing profiles. All other simulation conditions and parameters were set to average values (mouth size, temperature, oral volume, salivary flow rate), although these parameters are inherently individual. They could, in principle, be measured and used directly at the subject level. An alternative approach would be to represent these quantities as distributions rather than fixed mean values, and to sample from these distributions for simulation purposes. This would allow the generation of heterogeneous virtual panels and potentially improve the representativeness of the simulations compared to individual-level modeling.

### 4.4. Perspectives

#### 4.4.1. More products, more aromas, more consumption protocols behaviors

Future perspectives include extending the modeling framework to more complex product compositions, such as systems containing multiple aroma compounds as well as additional constituents (e.g., sugars, fats, and emulsifiers). In such systems, interactions between molecules could be explicitly accounted for, including partitioning effects, competitive binding, and matrix-dependent release mechanisms. This would allow a more realistic representation of structured food matrices, where aroma release is governed not only by interfacial mass transfer but also by the physicochemical properties of the surrounding components.

In addition, chemical processes occurring during oral processing could be incorporated into the model. This includes interactions with saliva, such as enzymatic activity, dilution, and potential metabolization or transformation of aroma compounds. Such mechanisms may significantly affect both the concentration and the temporal profile of volatile compounds in the oral and nasal cavities.

Overall, integrating these effects would provide a more comprehensive description of aroma release dynamics in realistic food systems, bridging physical transport phenomena with chemical and biochemical transformations.

#### 4.4.2. Considering 3D geometry of the oral cavity

Another promising direction is the integration of 3D models describing solid food breakdown during oral processing, with a stronger emphasis on physical mechanisms. Such approaches could rely on advanced simulation tools (e.g., SOFA, finite element methods) to quantify structural changes during mastication and bolus formation. These developments could build upon previous work in biomechanical modeling and material mechanics, including fracture of complex materials. The collaborative nature of the package enables researchers in the field to test and integrate their own models, thereby fostering collective progress toward a better understanding of the underlying mechanisms.

## 5. Conclusions

This study successfully demonstrated the development and validation of mechanistic *in vivo* models integrated into a modular Python-based framework for simulating aroma release during the consumption of liquid and melting solid foods. By representing the anatomical cavities as a series of interconnected reactors described by ordinary differential equations, the models effectively capture the dynamic transfer of volatile compounds from the food matrix to the nasal cavity. The integration of physiological parameters, such as salivary flow, bidirectional respiratory airflow, and discrete events like swallowing and mastication, proved essential for reproducing experimental observations. For solid matrices, the inclusion of temperature-dependent melting and dynamic surface area evolution allowed the model to accurately simulate the gradual release of aroma during oral processing. Validation against PTR-MS data from 30 subjects confirmed that the simulations could faithfully reproduce the primary kinetic features of retronasal aroma release for both aqueous solutions and gelatin gummy discs.

Looking forward, this flexible software package provides a foundation for exploring more complex food systems containing sugars, fats, and multiple aroma compounds. Future perspectives include the generation of heterogeneous virtual panels to account for inter-individual physiological variability and the integration of 3D biomechanical models to simulate the physical breakdown of solid food structures in the mouth. Ultimately, this tool offers a robust platform for the scientific community to further bridge the gap between food structure, oral processing, and sensory perception.

## Aknowledgements

The research leading to these results has received funding under grant (24.7 DIGIMOUTH) from Carnot Qualiment© (DOI: 10.17180/h5gd-gk88) supported by Agence Nationale de la Recherche (20 CARN 0026). This work was performed thanks to CALIS network’s resources (doi: 10.15454/9PGN-W156).

